# Effect of carbon nanotubes in electroactive neuron and cardiomyocyte differentiation on conductive 3D printed scaffolds

**DOI:** 10.64898/2026.01.08.698371

**Authors:** Lara Rodríguez, Goretti Arias, Gisela C. Luque, Maurizio Prato, Nuria Alegret

**Affiliations:** Carbon Bionanotechnology Group, CICbiomaGUNE, Basque Research and Technology Alliance (BRTA), Paseo Miramón 182, 20014 Donostia-San Sebastián, Guipúzcoa, Spain; Cardiac Diseases Group, Biogipuzkoa Health Research Institute, 20014 Donostia/San Sebastián, Spain; Polymer Reaction Engineering Group, INTEC (Universidad Nacional del Litoral-CONICET), Güemes 3450, Santa Fe, 3000, Argentina; Ikerbasque, Basque Foundation for Science, Bilbao, 48013, Spain; Department of Chemical and Pharmaceutical Sciences, Universitá Degli Studi di Trieste, Trieste, 34127, Italy

**Keywords:** Carbon nanotubes, 3D-printing, scaffolds, hiPSCs, neurons, cardiomyocytes

## Abstract

Carbon nanotubes (CNTs) have shown great potential in tissue engineering applications due to their unique properties, namely by improving electrical and mechanical properties of scaffolds. In recent years the use of 3D patterns, specially honeycomb or hexagonal patterns, to improve cell culture environment has also emerged in the tissue engineering field. Here we design HEMA-PEGDA based 3D printable scaffolds with and without CNTs in order to study the effect of both surface pattern and CNT incorporation on electroactive hiPSC-derived neuron and cardiomyocyte differentiation. Firstly, we tested scaffold biocompatibility with the SH-SY5Y neuroblastoma model, observing great viability and scaffold coverage for the CNT-containing formulation. As for the hiPSC differentiation models, we employed calcium signalling, immunocytochemistry and RT-qPCR techniques for cellular characterization. We found that CNTs and surface topography greatly affect neuronal culture maturation, by improving neuronal marker expression, calcium transient amplitude and axonal network maturation, while cardiomyocyte culture was mainly impacted by CNT presence independently of surface structure, although these conditions were not enough to reach full maturity. Overall, this study provided insights into the impact of surface structure and composition in electroactive cell differentiation and maturation.

## Introduction

In the tissue engineering field, creating *in vitro* models that closely resemble native tissue architecture and properties has become one of the main focus points during the last decade, as it allows for providing new insights into tissue function, disease modelling and drug development applications^1^ by filling the gap between 2D cell cultures and *in vivo* models^2^. To fulfil this purpose, it is essential to mimic the cellular microenvironment as closely as possible. Tissue specific microenvironment provides cells with cues to guide their cellular functions. Concentration gradients of soluble factors or contact with extracellular matrix components can modulate cell fate, especially in critical points such as developmental stages^3,4^, a complexity degree that is missing in 2D cultures^5^. This microenvironment is highly complex and difficult to mimic *in vitro*, different strategies having been developed to try and simulate native conditions as much as possible.

It has been reported for example that some cell types, such as neurons, respond to topographical features of their environment^6^, which has sparked the interest of modelling micro or nanometer scale structures and patterns to try and guide neuronal development^6,7^. Electrospinning, 3D-printed scaffolds and 2-photon lithography have been some of the techniques explored to create such structures^6–10^. Among them 3D printing is an interesting technique, as it allows to produce finely detailed structures with a plethora of tunable materials. For instance, scaffolds with hexagonal or honeycomb-like anisotropic patterns have been shown to promote cellular attachment and development in bone, neuron, cardiac and retinal cultures among others^11–15^, showing the potential in using architectural cues to promote 3D cellular interactions and network formation. Huang et al. developed a bilayer gelatin nanofiber scaffold with a honeycomb microframe in which they cultured and differentiated neurons and astrocytes. They found that cells formed clusters inside the honeycomb compartments and generated extremely intricate and interconnected 3D functional networks^16^.

Aside from controlling topography and microarchitecture, the application of nanomaterials and nanoparticles is also currently being extensively explored in the tissue engineering field. Carbon nanotubes (CNTs) for example, carbon nanostructures based on sp^2^-hybridized carbon atoms with a cylindrical shape^17^, have been the subject of multiple *in vitro* and *in vivo* studies for biomedical applications^18–21^. CNTs, in particular multi-walled CNTs, have been shown not only to improve mechanical and electrical characteristics of materials they are embedded in, but have also been employed in applications such as cardiac repair, bone regeneration and tissue healing^19,22,23^. Ye et al. for example studied neurogenesis in CNT-PEG hydrogel composites, describing optimal neuronal differentiation, marker expression and increased dendritic spine density for CNT composites^24^. Regarding cardiac tissue, Martinelli et al. designed PDMS-CNT scaffolds for cardiac repair. They studied neonatal rat ventricular myocyte (NRVM) cultures and found that in the presence of CNTs cells developed a more mature sarcomeric phenotype, increase of gap junction areas, electrophysiological phenotype and calcium transients, thus improving the overall maturation of cardiac myocytes^25^. Additionally, PEDOT/CNT substrates also had a significant impact of NRVMs, as CNT-substrates promoted homogeneous and higher beating amplitudes and mature phenotypical features like aligned sarcomeres and abundant Connexin-43 expression^26^.

Besides their unique properties, CNTs have been described to closely interact with cell membranes^27^, and even mechanisms by which CNTs promote neuronal growth and development having been explored. In particular, Shao et al. proposed a mechanism by which CNTs interacted with neuron membranes via integrin-mediated pathways, which in turn triggered downstream signalling pathways regulating cell survival, neurite growth, electrophysiological maturation and synapse formation of stem cell-derived neurons^28^.

Due to their multiple unique properties and improvement of cell cultures, CNTs have also been explored as substrates to control and enhance stem cell differentiation^29,30^. One of the biggest limitations of stem cell use is the acquisition of these types of pluripotent cells, an issue that has been overcome by the generation of induced pluripotent stem cells (iPSCs). These cells are generated by reprogramming adult somatic cells of donors and have the advantage of proliferating and expanding *in vitro*, solving the limited supply issue of stem cells^31,32^.

Taking all of this into consideration, in this study we designed and 3D-printed scaffolds with a hexagonal honeycomb pattern or a flat surface, either CNT-containing or CNT-free, in order to evaluate the effects of the surface topography on cell cultures. For this, we first evaluated scaffold biocompatibility using SH-SY5Y, a neuroblastoma cell line. We then seeded hiPSCs on the four types of scaffolds (CNT-containing L2CNT Pattern or Flat, and CNT-free L2 Pattern or Flat) and carried out differentiation protocols to achieve neuron and cardiomyocyte phenotypes. To study the differences between the four scaffold types on the resulting differentiated cell phenotype, we conducted calcium imaging to determine the functional activity and immunocytochemical and reverse transcription quantitative polymerase chain reaction (RT-qPCR) to study the expression of maturity markers. The results helped evaluate the effects both CNTs and surface patterns provide in differentiation of two types of electroactive cell cultures.

## Experimental Section

### Materials and 3D scaffold preparation

Carbon nanotubes (MWCNT, >95%) were purchased from Nanoamor Inc. (stock# 1237YJS: inner diameter, 5−10 nm; outside diameter, 20−30 nm; length, 0.5−2 μm). Water was purified with an Elix 3 system (Millipore, Molsheim, France). Hydrochloric acid (HCl, 37%), and 2-propanol were purchased from Sigma-Aldrich. 2-Hydroxyethyl methacrylate (HEMA), poly(ethylene glycol) diacrylate (PEGDA, average Mn 700 and Mn 250), and the photoinitiators, diphenyl (2,4,6-trimethylbenzoyl) phosphine oxide (TPO, Mn = 348.37 g mol−1) and lithium phenyl-2,4,6 tri-methylbenzoylphosphinate (LAP, ≥95 %) were all purchased from Sigma-Aldrich and used as received without further purification.

### CNT Purification

The initial step involved dispersing the CNTs (1 mg/mL) in a mixture of MilliQ water and HCl (1:1) through sonication for one hour. The resulting dispersion was then subjected to continuous stirring at 300 rpm overnight at room temperature. Subsequently, the CNTs were rinsed with MilliQ water until a neutral pH was achieved, followed by vacuum drying for 12 hours.

### Preparation and 3D printing of polymeric scaffolds

The hydrogel formulation for the scaffolds consisted of HEMA as the main component (monomer), TPO or LAP as the photoinitiator (1 wt.%), and 3 wt.% of PEGDA with two different molecular weights (Mn 250 or Mn 700) as the crosslinker. After combining the HEMA, crosslinker (PEGDA250 or PEGDA700), and photoinitiator (TPO or LAP), the formulation was sonicated for approximately 1 hour until a homogeneous ink was obtained and finally homogenized by using a vortex mixer (Heidolph instruments) for 1 minute at 2500 rpm. Additionally, 1 wt.% CNT was incorporated into the hydrogel formulation based on LAP and PEGDA250, requiring 2 hours of sonication. The composite formulations are presented in Table 1. The formulations were printed either in a honeycomb pattern or flat surface using an unmodified 3D printer (Photon mono), with adjustments made according to the printer’s and ink’s technical requirements. The light intensity of the 3D printer was set at 2 mW cm^−2^, and the exposure time per layer was adjusted for each formulation (Table 1). The initial layers, referred to as the bottom exposure, were subjected to a longer exposure time to ensure proper adhesion to the metal platform. After thickness (z) was set at 0.05 mm. After printing, the samples were soaked in isopropanol for 5 minutes, followed by a 5-minute water soak to remove any uncured resin (this process was repeated at least 3 times). Subsequently, the samples were washed in water three times and kept in water until further use.

**Table 1.**
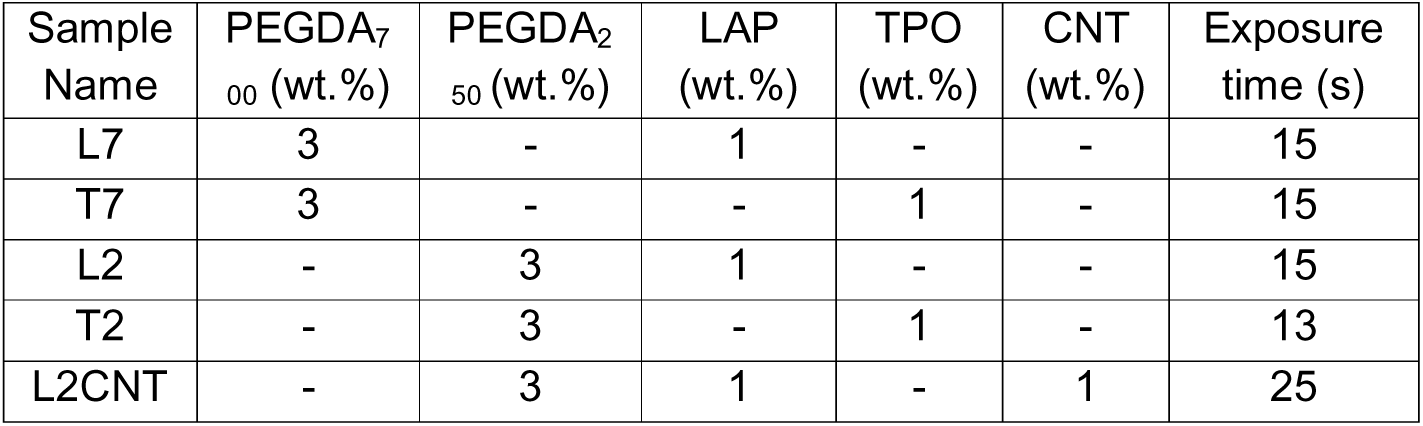
Formulations of inks for 3D-DLP printing. The first column presents the abbreviated names utilized in the subsequent discussion.

### Characterization of the polymeric scaffolds

#### Conductivity measurements

The conductive properties of the hydrogels containing CNTs were evaluated using sheet resistivity measurements. The sheet resistance (Rs) was evaluated using a conventional four-point probe method (Ossila, Sheffield, United Kingdom). The target current was set in a range between 10 and 100 μA with 10 V as the maximum voltage. The 3D printed hydrogels were kept in water overnight. Afterwards, the final thickness (t) of the wet films was measured with a calliper before the measurement was carried out. Conductive values reported for each polymer formulation is an average of at least three different samples. The electrical conductivity (σ) of the samples was calculated according to the following equation:

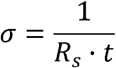

#### Tensile test

Mechanical properties were analyzed by performing tensile test in a universal testing machine (INSTRON 68SC-2) at room temperature. For tensile tests, films specimens polymerized with dumbbell shape of length 4 mm and cross section 0.49 mm. Assays were carried out with an elongation rate of 25 mm/min. Three samples were analyzed for each formulation.

#### Fourier Transform Infrared spectroscopy

To follow the polymerization, Fourier Transform Infrared spectroscopy (FTIR) was employed. Spectra were collected at various polymerization time points. Each spectrum corresponded to the average of 64 scans acquired within the 4000 -400 cm^-1^ spectral range, using a resolution of 4 cm^-1^. The absorption band at 1634 cm_⁻_¹ as was follows a function of polymerization time. FTIR measurements were performed using an Invenio-X spectrometer (Bruker), and the resulting spectra were subjected to baseline correction and peak analysis with OPUS vibrational spectroscopy software (Bruker).

#### Swelling

The swelling behavior of the scaffolds was analyzed in PBS solution at 37°C. At different times scaffolds were removed from PBS and weighed. Swelling ratio (S) was calculated by using the following equation: S = [(Wt−W0)/W0], where W0 is the weight of dry sample at time 0 and Wt is the weight of the swollen sample at time t.

#### Scanning Electron Microscopy Imaging

Scanning electron microscopy (SEM) images were obtained using a field-emission scanning electron microscope (FE-SEM; JSM-IT800HL, JEOL, Tokyo, Japan). Micrographs were recorded using an accelerating voltage of 2.5 kV and a working distance of 4 mm. For surface morphology evaluation, hydrogel samples were initially immersed in phosphate-buffered saline (PBS), plunge-frozen and subsequently freeze-dried. The dried samples were then mounted onto aluminum SEM stubs using double-sided carbon adhesive tape prior to SEM observation. To assess the internal architecture of the hydrogels, cross-sectional samples were immersed in PBS and sectioned using an ultramicrotome (Leica EM UC7) equipped with a cryo-chamber (Leica EM FC7). One side of each sample was trimmed with a scalpel at −70 °C. The resulting sections were freeze-dried, positioned with the cross-sectional face oriented upward, and mounted on aluminum SEM stubs using carbon adhesive tape. Imaging was carried out under the same SEM conditions described above.

Quantitative pore area analysis was performed using ImageJ software (National Institutes of Health, USA). Individual pores were manually segmented and measured, taking into account their specific morphology as observed in the SEM micrographs. To gain deeper insight into cell-surface interactions, cell-containing L2CNT scaffolds were further fixed with 2% Osmium tetroxide in PBS (Sigma-Aldrich) for 1 hour in the dark at 0°C. After thorough PBS washes, the scaffolds were plunge frozen in liquid nitrogen and freeze-dried for 24 hours. SEM images of cell seeded scaffolds were obtained with a JSM-IT800HL (JEOL, Japan) using 2.5 kV under high vacuum in secondary electron detection mode.

#### Thermogravimetric Analysis (TGA)

Thermogravimetric analyses were performed under air (25 mL·min^−1^ flow rate) using a TGA Discovery (TA Instruments). The samples were equilibrated at 100 °C for 20 min and then heated at a rate of 10 °C·min^−1^ in the range from 100 to 800 °C.

### In vitro studies

#### Neuroblastoma cell line culture

Neuroblastoma SH-SY5Y cells (AATC, CRL-2266) were cultured in complete DMEM:F-12 + GlutaMAX medium supplemented with 10% Fetal Bovine Serum and 1% Penicillin/Streptomycin at 37 °C and 5% CO_2_ in tissue culture-treated 75 cm^2^ flasks. Cells were passaged by incubating them for 5 min at 37 °C with 1X trypsin−EDTA solution. The detached cells were centrifuged at 1500 rpm for 5 min at 21 °C. The obtained pellet was then resuspended in 1 mL of complete DMEM medium. For cell counting, 10 µL of the cell suspension were diluted 1:2 in the exclusion dye Trypan Blue solution and counted with an automated cell counter (Countess, Invitrogen). Before cell seeding, the scaffolds were sterilized by autoclaving them and transferred to 8-well glass coverslips (Ibidi). Finally, media containing 500,000 SH-SY5Y cells was added to each well and incubated for 7 or 14 days at 37 °C in a humidified atmosphere with 5% CO_2_.

#### Lactate Dehydrogenase (LDH) Cell Viability Assay

The viability of cells adhered to the scaffolds was evaluated with a modified LDH CytoTOX96 Non-Radioactive Cytotoxicity Assay kit (Promega) reported by Ali-Boucetta et al^33^. Cell-containing scaffolds were mechanically disrupted by smashing after supernatant removal and addition of 150 μL of PBS containing 9% Triton X-100 (lysis buffer, LB). The samples were then frozen at −80 °C for 30 min, defrosted at 37 °C for 20 min, and smashed again. The cell debris was pelleted by centrifugation at 1000g for 10 min at 4 °C. After that, 50 μL of supernatants was transferred to replicate wells in a 96-well flat bottom plate. For LDH detection, 50 μL of substrate mix was added to the supernatants and incubated in the dark for 4 min, after which of 50 μL of stop solution were added. The absorbance at 492 nm was measured in a microplate spectrophotometer (BioTek Cytation 5, Agilent). LB alone was used as a negative control. All the collected data is represented as means of quadruplicates ± SD of three independent experiments (n= 3) and normalized to the average value of control sample L2.

#### Confocal imaging

Live/Dead staining was performed to evaluate cell viability using fluorescence imaging (LIVE/DEAD Cell Imaging Kit, Invitrogen). The reagent was prepared following the provider’s instructions. 24 hours after seeding the cells, the supernatants were discarded and replaced with new complete DMEM medium and Live/Dead reagent in a 1:1 ratio and incubated for 20 minutes at 37 °C and 5% CO_2_ before imaging. After 7 or 14 days of culture, the cells were fixed with 4% paraformaldehyde for 2 hours and stained with a dye against F-actin (ActinGreen 488 Ready Probes reagent, Invitrogen) in a 1:10 PBS dilution and DAPI (5µg/ml in PBS, Invitrogen) overnight at 4°C in the dark. The cell seeded scaffolds were then placed in 55 mm #1.5 coverslip glass bottom dishes (Cellvis). All the images were acquired in a laser scanning microscope (LSM 900 Axio Observer Z1, Zeiss). For the Live/Dead staining, 15 μm Z-stacks were acquired with the Plan-Apochromat 10x/0.45 M27 objective lens employing the excitation/emission wavelengths of 494 nm/400-567 nm for the Live (CalceinAM) probe and 570 nm/567-700 nm for the Dead (BOBO3) probe. For imaging the fixed scaffolds, 25 μm Z-stack images were generated with the same objective lens, employing the wavelengths of 353 nm/400−493 nm for DAPI and 493 nm/496-700 nm for ActinGreen 488. Orthogonal projection images based in the acquired Z-stacks were generated using ZEN 3.7 software. For quantification, fluorescence images were analyzed using the ImageJ program (National Institutes of Health, USA) by calculating the area taken up by cells in µm^2^ by using the 488 channel data in relation to the entire surface area for each scaffold. All the collected data is represented as means of triplicates ± SD of three independent experiments (n=3).

#### Human Induced Pluripotent Stem Cell (hiPSC) Culture

For cell differentiation studies, two hiPSC cell lines were utilized. For neuron differentiation, hiPSCs were purchased from The Jackson Laboratory-USA (JIPSC1100) and transduced with the pLVX-UbC-rtTA-Ngn2:2A:Ascl1 (pUNA) plasmid (Addgene, Plasmid #127289, MOI of 12), produced by the Viral Vectors Unit of the Spanish National Center for Cardiovascular Research (CNIC), Madrid, Spain. After 30 hours of incubating the hiPSCs in mTeSR Plus medium (StemCell) with polybrene and lentiviral particles, the media was replaced with fresh mTeSR Plus medium supplemented with 0.5 μg/ml of puromycin for the selection of transduced cells. For cardiomyocyte differentiation, hiPSCs from WiCell Research Institute-USA, were purchased (UCSD107i-2-6), a hiPSC cell line originated from skin fibroblast of a 74-year-old female donor without reported diseases. The cells were maintained in mTeSR Plus medium. For both types of hiPSCs, Matrigel (Corning) diluted in mTeSR Plus (20mg Matrigel/ml media) was used to coat culture plate surfaces prior to cell seeding to ensure attachment. Cells were passaged at least once a week by incubating with ReLeSR (StemCell) for culture expansion or Accutase (StemCell) for cell counting. After cell seeding, cultures were maintained with 10 µM Y-27632 RHO/ROCK pathway inhibitor (StemCell) in mTeSR Plus for 24 hours, after which the media was changed to fresh mTeSR medium. Cultures were maintained at 37°C with 5% CO_2_.

#### hiPSC differentiation into neurons

Autoclave-sterilized scaffolds were coated with Geltrex (1:97 dilution in mTeSR medium, ThermoFisher) and incubated overnight at 37°C. The coating was then removed and 250,000 hiPSC cells were seeded per scaffold and maintained with 10 µM Y-27632 inhibitor in the mTeSR medium. Following the protocol by Gantner et al^34^, the neuron differentiation process started 24 hours after cell seeding, by replacing the mTeSR medium with neuron conversion medium (Day 0). Briefly, the neuron conversion medium consisted of 1:1 DMEM/F-12 and Neurobasal Plus medium, supplemented with 2% B27, 1% N2, 1% GlutaMax, 1% Penicillin/Streptomycin, 1 μg/ml laminin, 2 μg/ml doxycycline, 10 μM SB431542, 200 nM LDN193189, 2.5 μM CHIR99021, 100ng/ml SHH C25II and 2 μM Purmorphamine. On day 13 the media was refreshed with neuron maturation medium, consisting of 1:1 DMEM/F-12 and Neurobasal Plus medium, supplemented with 2% B27, 1% N2, 1% ITS-A, 1% GlutaMax, 0.5% Penicillin/Streptomycin, 20 ng/ml GDNF, 20 ng/ml BDNF, 100 μg/ml Dibutyryl cAMP, 200 nM Ascorbic acid, 1 ng/ml TGFβ3, 10 μM DAPT, 1 μg/ml laminin and 2 μg/ml doxycycline. The media was refreshed every day until day 35 of culture.

#### hiPSC differentiation into cardiomyocytes

Similarly to neuron differentiation, autoclave-sterilized scaffolds were coated with Matrigel (20mg Matrigel/ml media) and incubated overnight at 37°C. The following day the coating was removed and 300,000 hiPSC cells were seeded per scaffold and maintained with 10 µM Y-27632 inhibitor in the mTeSR medium for 24 hours. For cardiomyocyte generation, the STEMdiff Cardiomyocyte Differentiation and Maintenance Kit (StemCell) was used following manufacturer’s instructions. Briefly, the cells were cultured in A, B and C conversion medium until day 8, after which the maturation medium was implemented until day 22 of culture. The cardiomyocytes were expected to start beating between day 12-18 of differentiation.

#### Calcium Imaging

At day 35 of culture for neurons and day 22 for cardiomyocytes calcium imaging was performed to assess the electrical activity of the cells, by incubating the cultures with Fluo-8AM (AAT Bioquest), a cell-permeant calcium indicator, at 2.5 µM for 20 minutes at 37 °C. A Cell Axio Observer microscope paired with a FITC filter and incubation chamber (Zeiss) was used to record calcium transients of spontaneous activity. For neurons, 500 ms/frame rate time-lapse videos were captured, while a 100ms/frame rate was used for cardiomyocytes. The amplitude and frequency of the calcium signals were analyzed for triplicate experiments (n=3) by making multiple recordings for each sample. Quantification of calcium measurements were performed with ImageJ (National Institutes of Health, USA) and reported as the fractional change in fluorescence intensity relative to baseline (F/F0). Within the sequence of images for each time-lapse, multiple regions of interest (ROI) for areas with calcium transients were selected, together with background areas that would determine the baseline fluorescence (F0). The temporal progression of fluorescence intensity values was determined for each ROI. At each time point, the fluorescence value was ratioed against the baseline value to yield F/F0. The amplitude value for each sample type was determined by averaging the maximum F/F0 values of transient peaks, while the frequency of the spikes was given by the time difference between adjacent peaks.

#### Immunocytochemistry

After calcium imaging, the cultures were fixed in 2% paraformaldehyde for 20 minutes, and further 60 minutes in 4% paraformaldehyde. Blocking solution (5% donkey serum, 2% BSA, 0.1% Triton X-100 and 0.1% NaN_3_ in PBS) was added after discarding the paraformaldehyde solution and incubated for 1 hour at room temperature. For neuron cultures, primary antibodies anti-NeuN (Millipore, ABN90) and anti-β-III-tubulin conjugated to Alexa Fluor 647 (Abcam, EP1569Y) were added and incubated at 4°C overnight, followed by the addition of secondary antibody Alexa Fluor 488 and DAPI and incubation at 4°C overnight. For cardiomyocyte culture, anti-sarcomeric _α_-actinin (Sigma-Aldrich, A7732) and anti-connexin-43 (Sigma-Aldrich, C6219) primary antibodies were utilized, following a 4°C overnight incubation, after which secondary antibodies Alexa Fluor 488 and 647 together with DAPI were incubated at 4°C overnight. Following the procedure employed for SH-SY5Y cell imaging, confocal images were acquired with a LSM 900 Axio Observer Z1 microscope, using excitation/emission wavelengths of 353 nm/400−493 nm for DAPI, 493 nm/496-700 nm for Alexa Fluor 488 (NeuN and _α_-actinin) and 570 nm/567-700 nm for Alexa Fluor 647 (β-III-Tubulin and Connexin-43). To allow quantification instrument settings were kept constant for all experiments.

#### Reverse Transcription Quantitative Polymerase Chain Reaction (RT-qPCR)

The RT-qPCR experiment was performed for neuron samples in order to further characterize the differentiation degree of the cultures. Total RNA content of all sample types, including additional 2D differentiation samples, was extracted using the miRNeasy Mini Kit and complementary RNase-free DNase kit (Qiagen) following manufacturer’s instructions. Extracted RNA concentration and purity was analyzed using a NanoDrop spectrophotometer. Reverse transcription to generate complementary DNA (cDNA) was performed using a SuperScript VILO cDNA synthesis kit (Invitrogen). The cDNA concentration and purity were also analyzed by the NanoDrop spectrophotometer. The RT-qPCR reaction was performed on a CFX384 Touch qPCR Detection System (Bio-Rad) with the SYBR Green system (Thermo Fisher). For a 384-well plate, with a final volume of 10 µl per well, comprised of 5 µl of SYBR Green master mix, 300 nM of each primer pair and 10 ng of cDNA were set up per well. Four biological replicas and three technical replicas were utilized per sample type, together with reverse transcription and no template negative controls. The following program was set for the analysis: 50°C for 2 min, 95°C for 10 mins, 95°C for 15s, 60°C for 1 min, repeating the third and fourth steps for 39 cycles. The analyzed genes were TUBB3, NGN2, ASCL1, SLC1 and SOX2, using GAPDH as the housekeeping gene. The utilized primer sequences are listed in Table S1 in the Supplementary Information. Relative expression quantification was performed using the 2^−ΔΔCt^ method, by normalizing the values to undifferentiated hiPSC control samples. Differences among groups were analyzed using a two-way analysis of variance (ANOVA), followed by Tukey’s post hoc test for multiple pairwise comparisons.

#### Statistical analysis

Statistical analyses for all experiments were performed using GraphPad Prism 9 software. For all the presented data except for RT-qPCR, differences among groups were analyzed using a one-way analysis of variance (ANOVA), followed by the Tukey’s post hoc test for multiple pairwise comparisons. Differences were considered statistically significant at *p<0.05, **p<0.01, ***p<0.001 and ****p<0.0001. All the reported data is presented as the means of triplicates ± SD of three independent experiments (n=3).

## Results and Discussion

### Mechanical properties

Mechanical uniaxial tensile tests to failure according to ISO 527 were performed on at least three dog-bone shaped specimens to evaluate the practical application of the 3D printed scaffolds. From the stress-strain curves (Fig. 1), the Young’s modulus (YM) was calculated.

**Figure 1.**
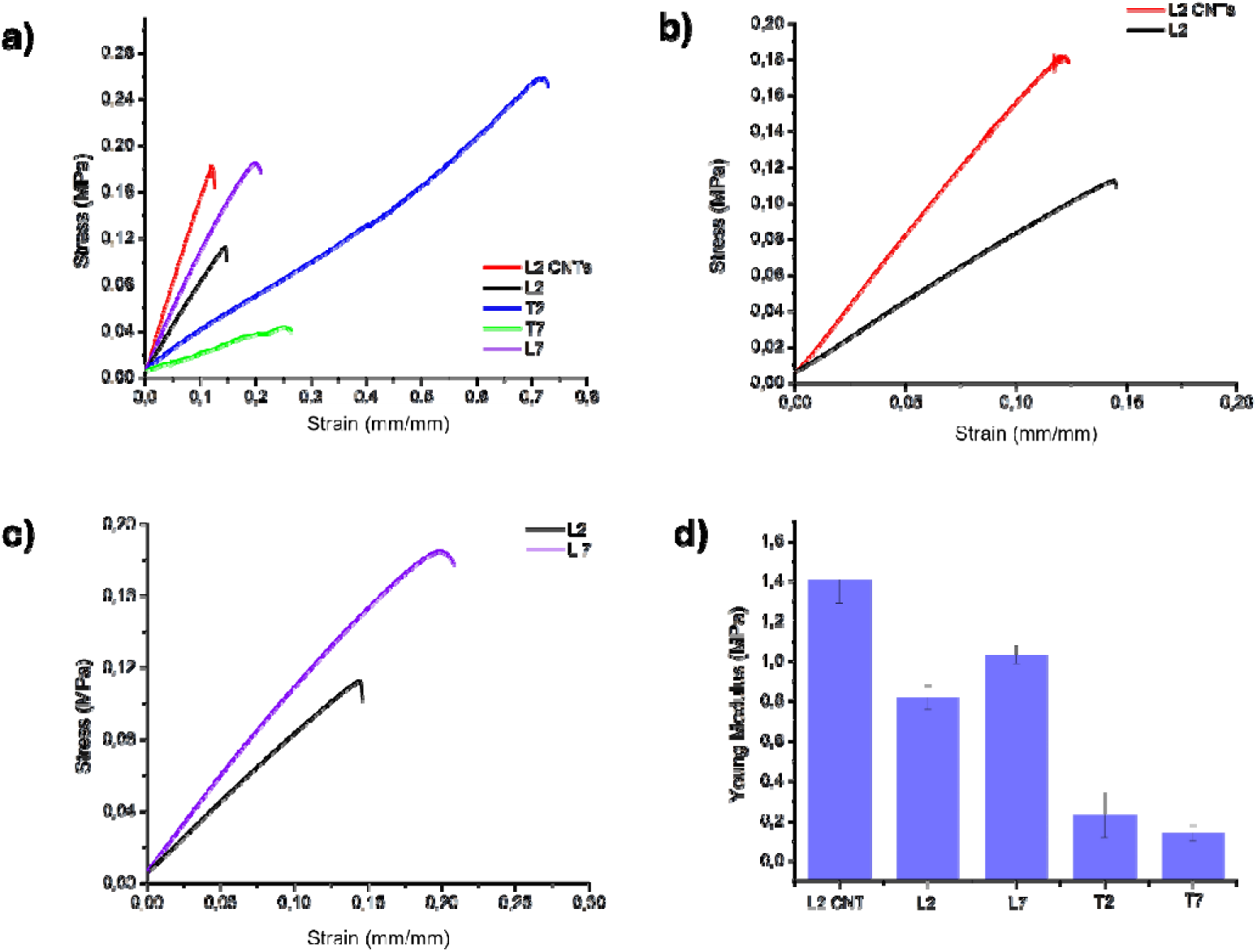
Stress-strain curves of the dog-bone shaped specimen hydrogels in the wet state. a) Comparation of different specimens, b) effect of CNT incorporation, c) effect of the photoinitiator, and d) Young’s modulus values.

From the mechanical tests, the effects of CNT incorporation, Mn, and the type of photoinitiator used in the formulations were analyzed. It can be observed that L2CNT presents a higher Young’s modulus (YM) of 1.4 MPa due to the incorporation of CNTs into the formulation, as expected and in agreement with previous reports in similar materials^35^. The Young’s modulus values for L7 and L2 are very similar, 1.03 and 0.82 MPa, respectively, suggesting that Mn does not significantly influence the mechanical properties of the final scaffolds. A similar behavior is observed for T2 and T7, with YM values of 0.23 and 0.14 MPa, respectively. When comparing the two photoinitiators, it is evident that the inks light-cured with LAP exhibit a higher YM than those photopolymerized with TPO, which can be attributed to the higher solubility of LAP in HEMA compared to TPO^36^, which leads to a higher amount of radicals generated and as a consequence a higher reticulation in the final scaffolds structure. Taking this into account and considering the good biocompatibility of LAP^37^, it was decided to further evaluate the effect of CNT incorporation in the formulation.

#### Fourier Transform Infrared Spectroscopy (FTIR)

To analyze scaffold polymerization after UV irradiation, Fourier transform infrared (FTIR) spectroscopy was performed. Figure S1 from Supplementary Information presents the FTIR spectra of L2 and L2CNT collected at different irradiation times. The absorption band at 1635 cm^-1^ corresponds to the carbon–carbon double bond of the HEMA monomer^38^ and was monitored to evaluate the progression of the polymerization process. As observed in the magnified FTIR spectra of both L2 and L2CNT, longer irradiation times were required to achieve polymerization in the presence of CNTs. This effect can be attributed to the ability of CNTs to absorb and scatter UV light, thereby reducing the effective irradiation reaching the polymer matrix and necessitating extended exposure times to obtain a comparable degree of polymerization.

#### Electrical conductivity

Electrical conductivity is a key parameter in materials designed for tissue engineering, especially for tissues composed of electroactive cells, as it enables the reproduction of native bioelectrical cues, directs cell growth (e.g., nerve, muscle, bone, and cardiac tissues), and supports functional tissue regeneration^39^.

In order to determine the electrical conductivity of the scaffolds containing CNTs, three samples were prepared, and three measures were done in each scaffold. Table 2 summarizes the results of the electrical conductivity for L2CNT scaffolds. M1 to M3 represent the average values obtained from three measurements performed at different locations on each scaffold. The measured conductivities are consistent with those previously reported for comparable CNT-containing materials. For instance, Ma et al.^40^ described an alginate gelatin-based scaffold containing 1 % wt. of CNTs, which exhibited electrical conductivity values of approximately 2 x 10-4 S cm^-1^.

**Table 2.** Electrical conductivity of the L2CNT scaffolds.

| L2CNT | M1 | M2 | M3 |
| --- | --- | --- | --- |
| Conductivity (S/cm) | $3,5 \times 10^{-4}$ | $3,9 \times 10^{-4}$ | $4,2 \times 10^{-4}$ |
| Error | $0,4 \times 10^{-5}$ | $0,5 \times 10^{-5}$ | $0,6 \times 10^{-5}$ |

Notably, the conductivity values were comparable across different regions of the same scaffold as well as among different samples, indicating a homogeneous distribution of CNTs throughout the polymeric scaffold.

#### Swelling

Swelling behavior is a key parameter in the characterization of polymeric scaffolds to be employed in biomedical applications. For *in vitro* tests, when scaffolds are exposed to physiological fluids, these can absorb and undergo volumetric changes that directly affect their porosity, pore interconnectivity, and mass transport properties. For cell adhesion, proliferation, nutrient diffusion it is important to have a scaffold able to provide nutrients trough the structure, but also to maintain the shape and mechanical properties^41^. Figure S2 in the Supplementary Information shows the swelling degree of the five scaffolds measured in PBS at 37 °C. The results clearly highlight the influence of the molecular weight (Mw) of the crosslinker on the swelling behavior of the scaffolds. Scaffolds crosslinked with PEGDA 700 exhibit the lowest swelling degree, reaching values of approximately 20% after 4 h of incubation. This behavior can be attributed to the formation of a denser polymeric network with reduced effective porosity, which limits PBS solution uptake and restricts network expansion. In contrast, scaffolds crosslinked with PEGDA 250 display higher swelling degrees, in the range of 40 - 50% at the same time point. The lower molecular weight of the crosslinker likely results in a less densely packed network and increased chain mobility, facilitating greater PBS absorption and volumetric expansion. Regarding the effect of CNT incorporation, the swelling behavior of CNT-containing scaffolds remains comparable to that of their CNT - free counterparts. No significant differences in swelling degree are observed after 4 h of immersion, indicating that, at the concentrations employed, CNTs do not substantially affect the hydrophilicity or crosslink density of the polymer network.

#### Scanning electron microscopy imaging (SEM)

For *in vitro* hiPSC differentiation on scaffolds, not only mechanical properties and electrical conductivity are important, but also scaffold morphology and porosity, as these features are critical for promoting cell adhesion and proliferation. Therefore, the surface morphology of the scaffolds was analyzed using scanning electron microscopy (SEM). SEM micrographs (Fig. 2) show that the scaffolds were successfully 3D printed in the honeycomb pattern, with honeycomb shaped pores in the 300 µm diameter range. Pore size is a well-established determinant of cell attachment, growth, and proliferation, with optimal pore diameters reported to range between 80 and 150 µm for neuronal cells^42^ and between 25 and 60 µm for cardiomyocytes^43^. To further investigate differences in porosity among the formulations selected for hiPSC differentiation into neurons and cardiomyocytes (L2 and L2CNT), scaffold cross-sections were prepared and examined by SEM, enabling visualization of the internal pore architecture beyond the surface layer. As shown in Figure S3 in the Supplementary Information, L2CNT scaffolds displayed a noticeably higher pore density per unit area compared with L2 scaffolds, with average pore sizes of approximately 50.9 µm and 22.4 µm, respectively. Moreover, as mentioned previously, L2CNT scaffolds exhibit a homogeneous distribution of CNTs, which is also evident in surface SEM images (Fig. 2 c). In these images, an increase in surface roughness and surface area is observed, which can be attributed to the incorporation and organization of carbon nanotubes within the polymer matrix during the polymerization process, making possible for cells cultured on the scaffolds to have direct contact with CNTs.

**Figure 2.**
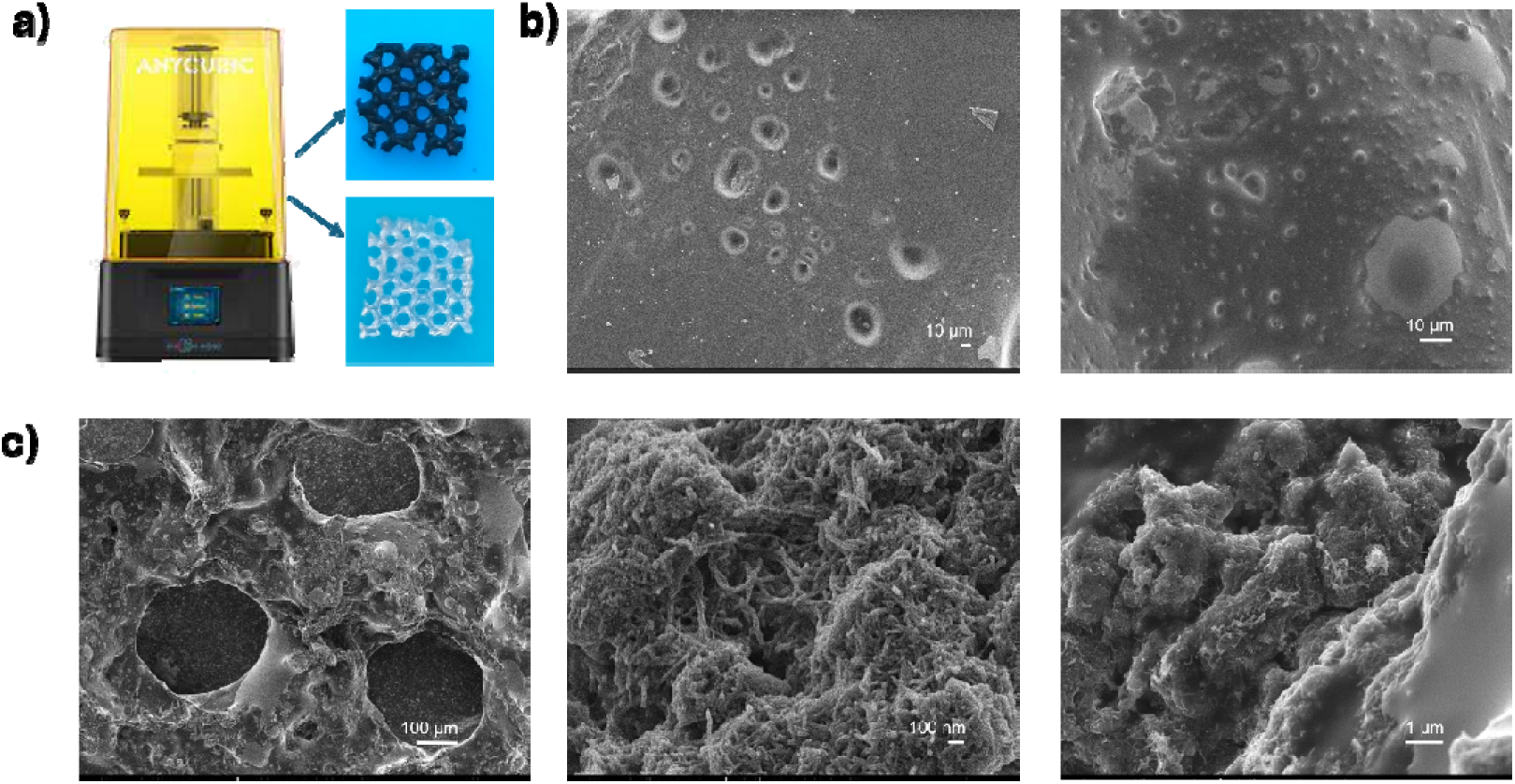
Surface L2 and L2CNT scaffold micrographs. a) Pictures of 3D printed scaffolds L2 and L2CNT, b) SEM images acquired at an accelerating voltage of 2.50_JkV of the surface of L2 c) and L2CNT.

#### Thermogravimetric analysis (TGA)

With the aim to analyze the thermal stability of the scaffolds selected for *in vitro* testing, TGA analysis for L2 and L2CNT scaffolds was performed (Fig. S4 in Supplementary Information). As it can be observed, both scaffolds are stable up to 200°C. Therefore, autoclaving at 121°C was chosen as the sterilization method prior to *in vitro* studies.

### Biocompatibility studies in the neuroblastoma SH-SY5Y cell line

In order to perform a first biocompatibility trial for the scaffolds, SH-SY5Y cells, a neuroblastoma model known for its neuron-like phenotype^44^, were seeded onto all the formulations (500.000 cells/scaffold). Live/Dead assays were carried out 24 hours after seeding, (Fig. S4 from Supporting Information), finding no differences among the groups, which suggests low short-term cytotoxicity and good viability for all formulations. The scaffolds were incubated for either 7 or 14 days *in vitro*, after which viability assays and immunocytochemical stainings were performed. Out of all the preliminary formulations (L2, L7, T2 and T7), the formulations prepared with the LAP photoinitiator demonstrated a higher viability rate compared to those with TPO, among which L2 presented the best values for both timepoints (Fig. 3 a and c) and was therefore selected as the formulation to further test with and without CNTs. The results for all the formulations are summarized in Figure 3. SH-SY5Y cells exhibited higher viability for CNT containing hydrogels (L2CNT) as opposed to its control counterparts (L2), after both 7 and 14 days of culture. After fixation, the samples were stained with ActinGreen, a commercial dye for cytoskeletal F-actin, and the nuclear dye DAPI. Confocal images of the whole scaffolds were acquired and analyzed by quantifying the area of the scaffold covered by cells compared to its whole surface area (%Coverage). The cell coverage percentage is significantly higher for L2CNT, for both 7- and 14-day samples, showing a 34% and a 35% increase compared to L2 respectively. As for the rest of the formulations, they all performed similarly in the absence of CNTs (ranging from 2-8% cell coverage). Some studies have described an increased amount of lamellipodia and cell-cell contacts in SH-SY5Y cells when culturing them on CNT containing PEDOT scaffolds compared to control conditions^45^, which hints at the ability of CNTs to enhance cellular adherence. Finally, in order to observe the cell-scaffold interactions, SEM imaging was performed to L2CNT scaffolds (Fig. 4). A cell monolayer covering the scaffold surface can be observed, confirming the high cell coverage results. Furthermore, areas where the cells have developed a neurite-network can also be observed, suggesting a more differentiated state of the SH-SY5Y cells. These results line up with previous studies showing that in the presence of CNTs SH-SY5Y cells exhibit not only higher proliferation rates, but also a greater differentiation degree. For instance, Yoon et al. described higher viability, neural marker expression levels and neural activity on SH-SY5Y cells when cultured on a CNT surface compared to control polystyrene and graphene films^46^. All together these findings confirm the non-cytotoxic nature of the scaffolds and the substantial positive impact of the CNT addition on SH-SY5Y cells, making these scaffolds suitable candidates to support and enhance hiPSC-derived electroactive cell differentiation.

**Figure 3.**
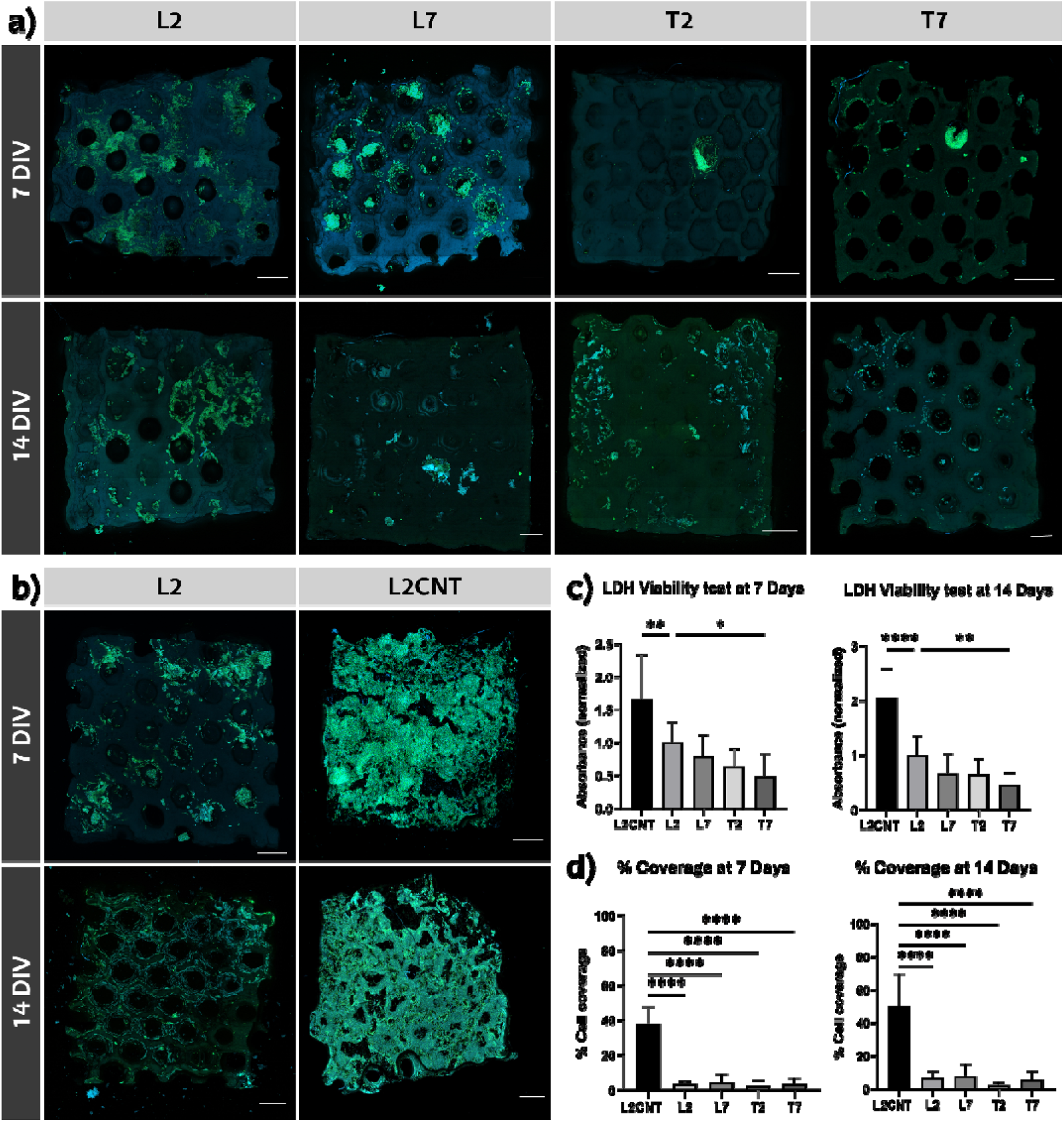
SH-SY5Y neuroblastoma cells cultured on the scaffolds at 7 and 14 days *in vitro*. a) Orthogonal projections of the confocal images taken for the L2, L7, T2 and T7 formulations, b) Orthogonal projections for the L2 control and L2CNT carbon nanotube containing scaffolds, c) LDH viability test results, d) Percentage of cell coverage of the scaffold surface. Cytoskeletal F-actin shown in green (ActinGreen, 488 nm) and nucleus in blue (DAPI, 405 nm). *p-value < 0.05, **p-value < 0.01, ***p-value < 0.001, ****p-value < 0.0001. Scale bars= 500 µm.

**Figure 4.**
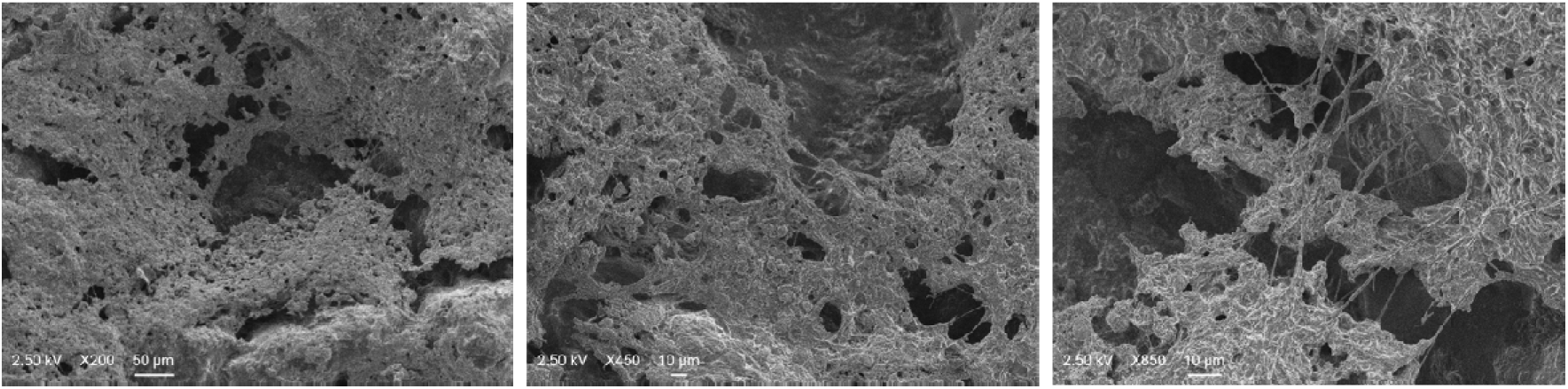
SEM images of the SH-SY5Y cells in the surface of the L2CNT scaffold.

### hiPSCs culture and differentiation

Many studies have shown the benefits of an electroconductive environment for cell differentiation, especially regarding electroactive cells. The incorporation of an electroconductive material, such as CNTs, has proven to enhance cellular differentiation and electrophysiological activity. Shin et al. have described improved calcium signalling for neural stem cells differentiated in hyaluronic acid-CNT scaffolds, by upregulating the expression levels of voltage-gated calcium channels^47^. CNT-multilayered scaffolds have also been described to regulate neuron maturation by integrin-mediated interactions, causing downstream regulation of cell survival, proliferation, differentiation and synapse formation mechanisms^48^. Not only neurons have been described to benefit from CNT interactions. Zhang et al. proposed a mechanism via NCX1 regulation, a sodium/calcium channel vital for calcium homeostasis in cardiomyocytes, by which CNTs promote contraction and functional activity in neonatal rat ventricular cardiomyocytes^49^.

Although a conductive environment is clearly beneficial for the development of certain types of cultures, multiple studies have also pointed to the positive effect of patterns in cell differentiation to adult phenotypes, as it is recognized that three-dimensional (3D) cultures more closely resemble *in vivo* conditions compared to traditional 2D cultures. For instance, when culturing and differentiating hiPSC cells on microstructured 3D honeycomb scaffolds, cells were described to generate intricate interconnected multilayer functional neuronal networks^9^. Among the wide variety of existing patterns, the honeycomb type designs have been widely utilized for tissue engineering applications due to their positive impact on network formation and cell maturation on multiple cell types^13,16,50,51^. In order to define the impact of the honeycomb pattern in cell differentiation, L2 and L2CNT scaffolds were also printed as a flat surface with no pattern (L2 Flat and L2CNT Flat from here onwards).

Taking all of this into consideration, we decided to test the effect of our CNT-containing scaffolds in electroactive cell cultures, specifically the effect on hiPSC differentiation into neurons and cardiomyocytes. For neuron differentiation, hiPSC cells transduced with a pUNA plasmid, containing neurogenesis regulator genes Ngn2 and Ascl1^52^, were utilized, while for cardiomyocyte conversion a commercially available WiCell hiPSC line was chosen.

Therefore, hiPSCs were seeded onto control (L2 Pattern and L2 Flat) and CNT-containing (L2CNT Pattern and L2CNT Flat) scaffolds. For obtaining neurons, the hiPSC-pUNA cells were cultured following a neuron differentiation protocol for 35 days *in vitro,* while for cardiomyocyte induction, the differentiation process for WiCell hiPSCs lasted 22 days following the StemCell Cardiomyocyte Differentiation kit.

### Calcium Imaging

To assess the functional activity of the cultures after the differentiation process ended, calcium imaging was performed by incubating the samples with Fluo-8AM reagent, a fluorescent dye that allows the visualization of calcium influxes in the cells. Spontaneous calcium transients were observed for both neuron and cardiomyocyte cultures (Video S1 and S2 respectively), pointing to a successful differentiation process for both phenotypes by confirming an inherent electrophysiological activity without external stimulation. The data obtained by calcium imaging was then analyzed to determine the signal amplitude and frequency (period) of each culture type (Fig. 5). Regarding neuronal cultures, calcium spikes showed the highest amplitudes in L2CNT Pattern samples, followed by both types of L2 scaffolds, L2CNT Flat presenting the lowest values among all. However, when looking into the frequency of the spikes, both L2 and L2CNT Pattern showed significantly lower values (10.2 and 13.3 s respectively) compared to Flat scaffolds, mainly L2 Flat (345.9 s), indicating a significantly faster spike frequency for neurons differentiated on Pattern surfaces. Overall, these results would point to a greater impact of the surface pattern on neuronal activity rather than the presence or absence of CNTs. As described by Koroleva et al, this outcome could be the consequence of neural networks reaching greater maturity, and thus, functionality, when cultured in an environment that promotes the generation of a multilayer network as opposed to a flat surface^9^. However further characterization in necessary to support such hypothesis. As for cardiomyocytes, while period values were in the same range (1-1.5 s) for all samples, L2CNT scaffolds, both Pattern and Flat, presented significantly higher amplitude values compared to both types of control L2 scaffolds. CNT enriched scaffolds have been reported to increase calcium oscillation homogeneity, frequency^25^, rhythm stability and fluctuation amplitude^53^ in neonatal rat ventricular cardiomyocytes. The significantly enhanced amplitude values highlight the positive influence of CNTs on the functionality of cardiomyocyte cultures, whereas we did not a quantifiable effect produced by the pattern of the scaffold surface.

**Figure 5.**
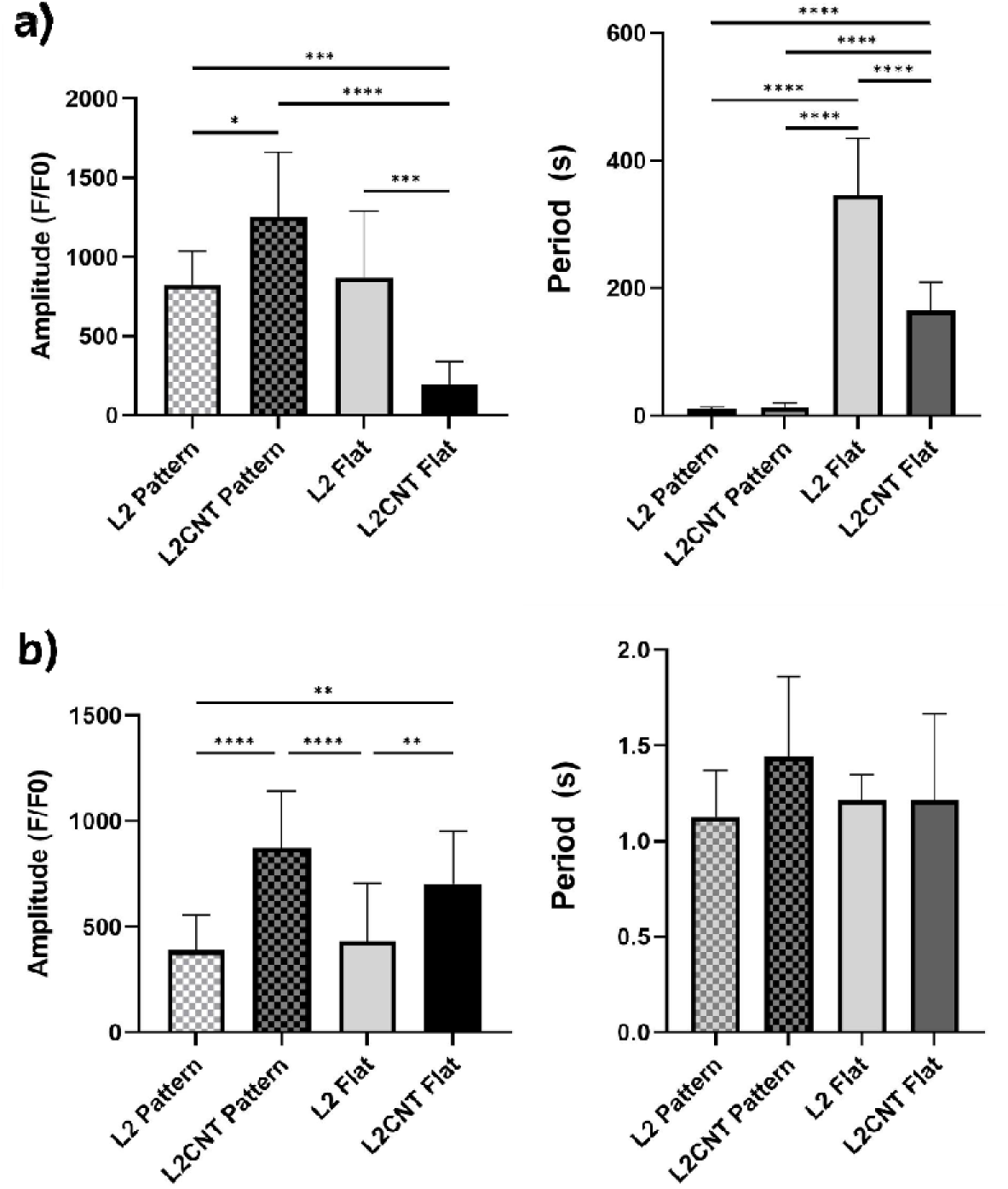
Calcium imaging results for differentiated neuron and cardiomyocytes cultured on the scaffold. a) Average amplitude and period values for differentiated neurons, b) Average amplitude and period values for differentiated cardiomyocytes. *p-value < 0.05, **p-value < 0.01, ***p-value < 0.001, ****p-value < 0.0001.

### Immunocytochemistry

To further characterize the phenotypical maturity of the obtained cultures, the samples were fixed to perform immunocytochemistry. For neuron cultures, the cells were stained against NeuN, a nuclear specific protein, and β-III-Tubulin, a microtubular component, both found in mature neurons and used as neuronal markers^54,55^. All sample types were positive for NeuN and presented similar intensity values for this marker (Fig.6 a and c), indicating a proper mature neuronal phenotype^56^ across all scaffold types. Interestingly, we found that neurons differentiated on Pattern scaffolds presented a greater number of long axons, especially in L2CNT Pattern, where the axons extend over the entire surface of the scaffolds and even over the 300 µm honeycomb compartments creating a network that connects both sides of the pores (Fig. 6 b). Neuronal clusters were observed mainly in the honeycomb pores interconnecting different clusters, similarly to reports by Huang et al^16^. This correlates to the significantly higher β-III-Tubulin intensity values observed for these samples (Fig. 6 c), followed by those of L2 Pattern. β-III-Tubulin is particularly relevant to ensure the growth and guidance of axons, reaching peak expression levels during neuronal maturation stages^56,57^. This result is therefore in line with the characteristics for neuronal activity observed for calcium imaging (Fig. 5), as these images together with calcium transient data support the hypothesis of a mature neural network found in pattern scaffolds, particularly for L2CNT.

**Figure 6.**
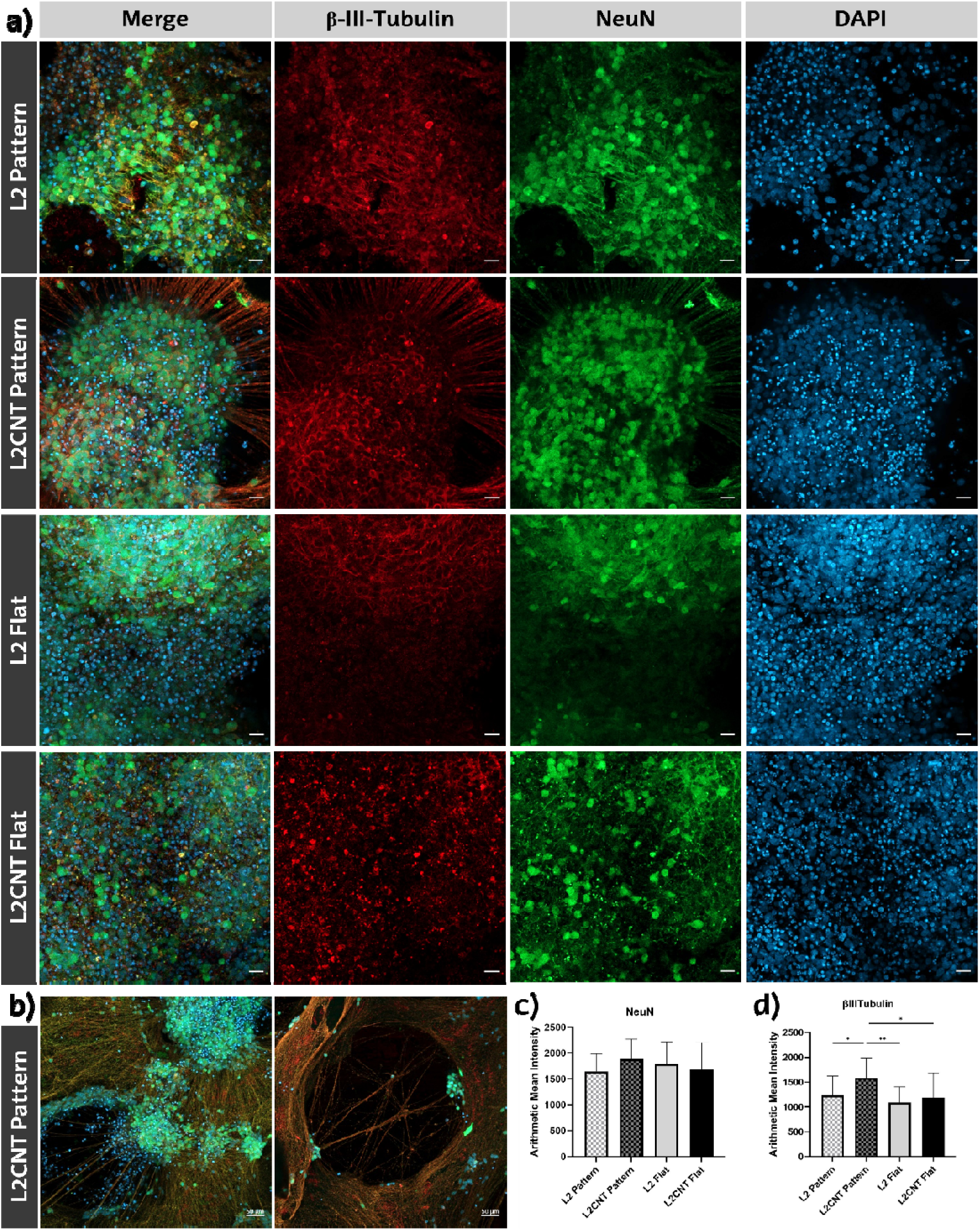
hiPSC-derived neurons differentiated on the scaffolds at 35 days *in vitro*. a) Orthogonal projections of the confocal images taken for flat and honeycomb-pattern L2 and L2CNT scaffolds at 20x (scale bars= 20 µm), b) Confocal images for L2CNT scaffolds at 10x where axon extension can be observed, c) Average arithmetic mean intensity values for Neun (488 nm), d) Average arithmetic mean intensity values for β-III-Tubulin (647 nm). Microtubular β-III-Tubulin shown in red (647 nm), nuclear NeuN protein shown in green (488 nm) and nucleus in blue (DAPI, 405 nm). *p-value < 0.05, **p-value < 0.01.

Although most cultured cells were positive for neuronal markers, and phenotypical characteristics of neurons like axons were present, we also observed a subpopulation of cells by DAPI staining that were negative for these markers (Fig.6 a), probably due to some degree of heterogeneity among the cellular population. Neuron subpopulation heterogeneity upon differentiation has been thoroughly studied by Lin et al. According to their single-cell RNA-sequencing data, neuron-induction via Ngn2 pathway, as in our study, produces a variety of heterogeneous neuronal populations with distinct molecular expression patterns^58^.

As for cardiomyocytes, the cells were stained against sarcomeric _α_-actinin, a key element in cardiac muscle sarcomeres, and Connexin-43, the main protein responsible for gap junction formation in cardiac muscle^59,60^. In contrast with neurons, a clear mature cardiomyocyte phenotype could not be confirmed for either formulation, as the _α_-actinin staining failed to define sarcomeric structure, either because of an unspecific immunostaining or a not fully mature phenotype. However, Connexin-43 did provide specific staining for cell junctions in the case of L2CNT formulations, which also showed bigger nucleus size and more organized cell distribution (Fig. 7) compared to L2 controls. Although calcium imaging confirmed the functionality of the cardiomyocyte culture, the immunocytochemical studies determine that CNT presence or a surface pattern is not sufficient for cells to achieve full phenotypical maturity in this type of scaffolds.

**Figure 7.**
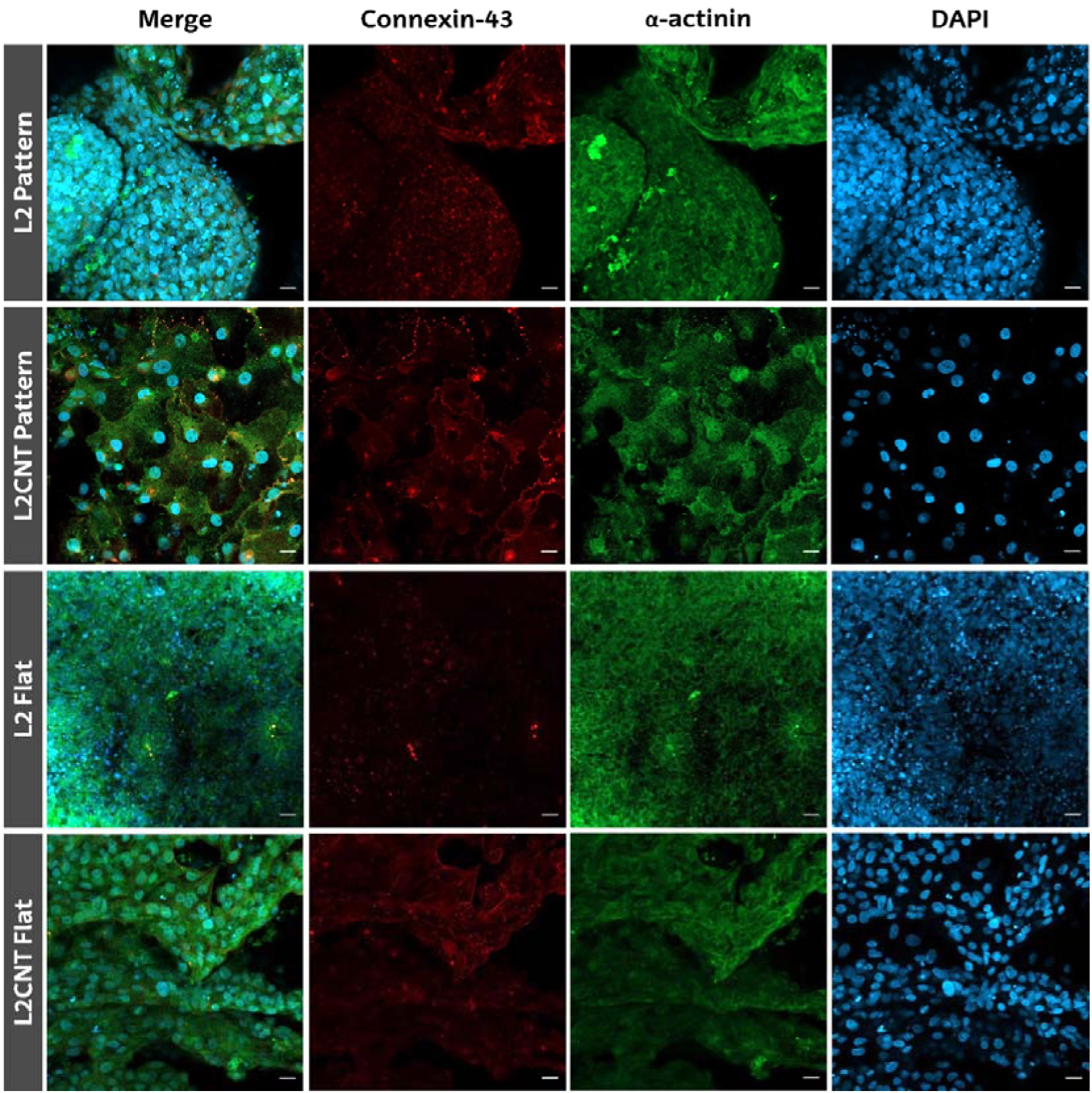
hiPSC-derived cardiomyocytes differentiated on the scaffolds at 22 days *in vitro*. Orthogonal projections of the confocal images taken for flat and honeycomb-pattern L2 and L2CNT scaffolds. Connexin-43 shown in red (647 nm), sarcomeric _α_-actinin shown in green (488 nm) and nucleus in blue (DAPI, 405 nm). Scale bars= 20 µm.

### Reverse Transcription-Quantitative Polymerase Chain Reaction (RT-qPCR)

As to further characterize the differentiation degree of the neuron cultures and make sense of the calcium imaging and immunocytochemical results, we decided to perform a RT-qPCR experiment to measure mRNA expression levels. We used undifferentiated hiPSC-pUNA cells as control samples and 2D-differentiated neurons to compare the effect of the scaffolds on the differentiation process. We analyzed SOX2 as a pluripotency marker^61^, Ngn2 and Ascl1 neurogenesis markers^62,63^, SLC1A1 a marker for a glutamate transporter present in neurons^64^, and TUBB3, the β-III-Tubulin neuronal marker^65^ (Fig. 8).

**Figure 8.**
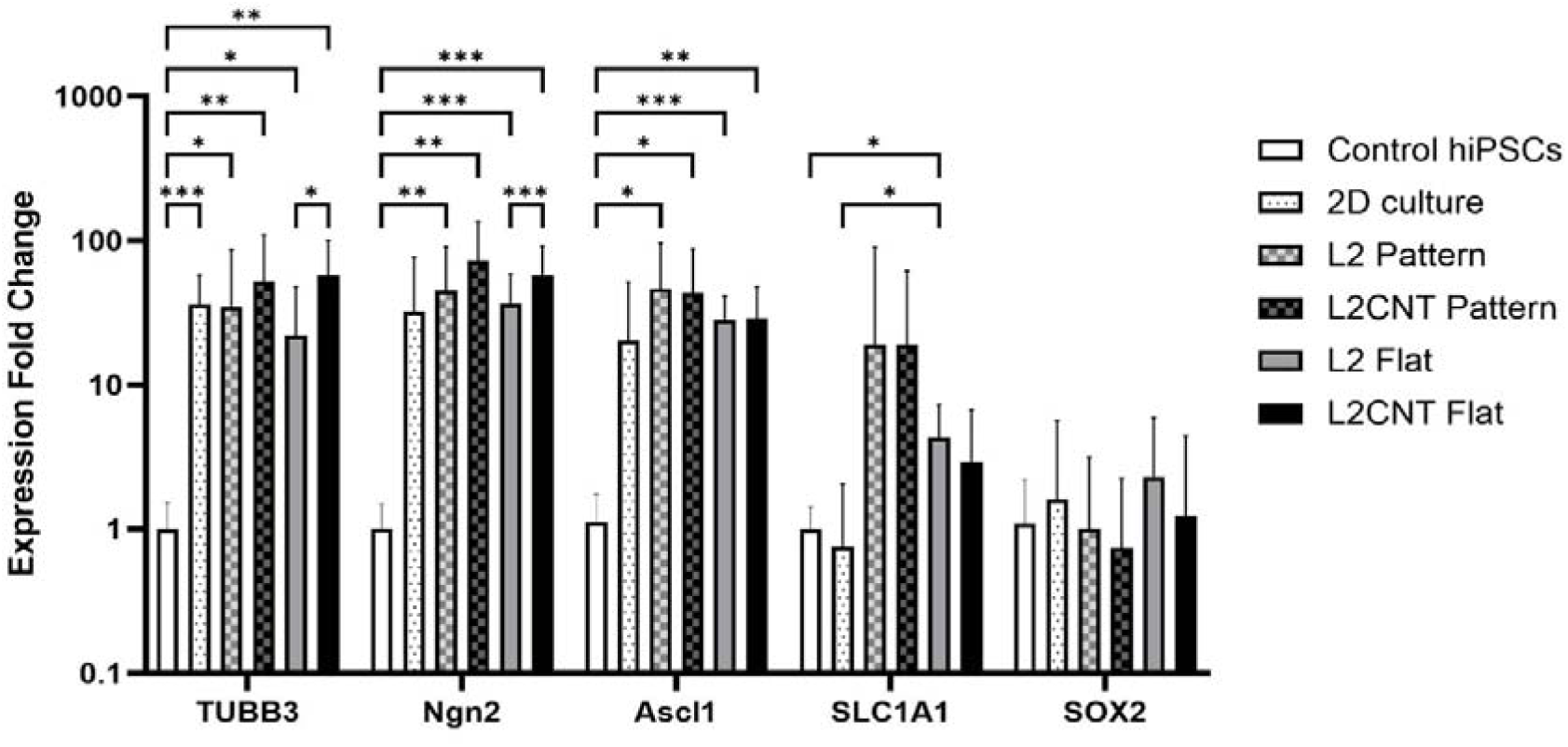
RT-qPCR results for differentiated neurons (n=4). Neuronal markers TUBB3, Ngn2, Ascl1, SCL1A1 and pluripotency marker SOX2 were analyzed. *p-value < 0.05, **p-value < 0.01, ***p-value < 0.001.

TUBB3, Ngn2 and Ascl1 expression levels were the highest across all types of surfaces compared to hiPSC-pUNA control. Among the different types of scaffolds studied, L2CNT, both Pattern and Flat, presented the greatest expression levels for TUBB3 (fold change= 52.5 and 56.9 respectively) and Ngn2 (fold change=72.5 and 57.2 respectively). Ascl1 expression however was predominantly influenced by the pattern of the scaffolds, appearing to have higher expression levels in Pattern L2 (fold change=46.5) and L2CNT (fold change=43.4) scaffolds compared to Flat ones (fold change=28.1 for L2 and 29.2 for L2CNT). Regarding SLC1A1, Pattern scaffolds were the only ones showing significantly higher expression levels compared to control (fold change=19.08 for L2 and 19.05 for L2CNT), followed by Flat scaffolds (fold change=4.3 for L2 and 2.9 for L2CNT). Interestingly, 2D cultured neurons presented the lowest expression levels together with L2 Flat for all neuronal markers. These results further evidence that CNT presence and surface pattern greatly improve the maturation degree of cultured neurons.

For SOX2 pluripotency marker expression, there were no significant differences between any of the samples, once more hinting at the presence of more than one neuronal subpopulation^58^. Although SOX2 is generally regarded as a pluripotency marker, for some differentiated neurons and glial cells SOX2 expression is essential for normal development and function^66,67^. Therefore, it is possible for a subpopulation of our cultures to be responsible for the observed SOX2 expression, being either SOX2 positive neuronal cells or residual hiPSCs.

Overall, the RT-qPCR results strengthen the observations raised from previous experiments, in that the L2CNT formulation enhances neuronal differentiation into an electroactive mature phenotype when structured in a honeycomb pattern.

## Conclusion

In the present study CNT-containing HEMA-PEGDA scaffolds were successfully prepared and 3D printed into honeycomb and flat designs. Four formulations were preliminary designed, consisting of a HEMA matrix and combinations of PEGDA_250_ or PEGDA_700_ crosslinker and LAP or TPO photoinitiators. Out of all formulations, after mechanical testing, L2, containing PEGDA_250_ and LAP photoinitiator, was selected due to higher values observed for its Young’s modulus and reported LAP biocompatibility^37^. L2 and its CNT-containing counterpart, L2CNT, which presented desirable and homogeneous conductivity values, were further studied before *in vitro* testing. SEM analysis confirmed proper 3D printing of the scaffolds in a honeycomb pattern and revealed bigger average pore sizes in the internal structure of L2CNT (50.9 µm) compared to L2 (22.4 µm), with CNTs being observed on the surface, accessible for cell-CNT interactions.

When testing these CNT scaffolds as growth substrates for SH-SY5Y cells, they proved to be highly biocompatible and improve cell adhesion compared to L2 control substrates. After the first set of successful *in vitro* tests with a neuroblastoma cell line, it was decided to evaluate the effect of CNT presence and surface pattern on the differentiation of electroactive cells, in particular neurons and cardiomyocytes, by calcium imaging, immunocytochemistry and RT-qPCR analysis. Regarding hiPSC-derived neuronal cultures, it was determined that the 3D honeycomb pattern greatly improved neuron maturation by creating 3D interconnected axonal networks with calcium spikes faster and greater in amplitude, cells positive for NeuN and β-III-Tubulin and significantly higher neuronal marker expression levels (TUBB3, Ngn2, Ascl1 and SLC1A1). These effects were accentuated in the presence of CNTs in the scaffold, indicating that the combination of a 3D pattern and the conductive properties of the CNTs are extremely beneficial for neuronal cell differentiation and phenotypical maturation. As for hiPSC-derived cardiomyocytes, functional maturity was observed when studying calcium transients, with homogeneous beating, the amplitude of the calcium transients being enhanced by CNTs. However, although functional activity was defined, phenotypical maturity could not be confirmed, as Connexin-43 staining was successful for L2CNT scaffolds, but not sarcomeric _α_-actinin staining. Thus, although CNTs seem to improve cardiomyocyte activity independently of the surface pattern, these conditions are not sufficient for these types of cells to reach full maturity.

Overall, we conclude that in this study we managed to create and 3D print CNT-containing patterned scaffolds which demonstrated high biocompatibility and improved electroactive cell culture maturity, especially regarding neurons.

## Author contributions

LR conducted the scaffold fabrication and *in vitro* experiments, performed statistical analyses and wrote the manuscript (Investigation, Formal analysis, Writing – original draft). GL conducted the material characterization experiments and contributed to writing the manuscript (Investigation, Writing – original draft). GA and NA conceived the study (Conceptualization). NA provided critical revisions and supervised the work (Writing - review & editing, Supervision). MP supervised the project and secured funding (Supervision, Resources, Funding acquisition).

## Data availability

The data supporting this article have been included as part of the Supplementary Information. Supplementary information: Table S1 and further experimental details.

## Acknowledgements

This research was supported by the Center for Cooperative Research in Biomaterials (CIC biomaGUNE) and the Biogipuzkoa Health Research Institute (Biogipuzkoa HRI). It was carried out under the Spanish Research Agency (PID2022-140419OB-I00), the IKUR Strategy 2022 (NANONEURO) on behalf of the Department of Science, Universities and Innovation of the Basque Government, and received additional funding from the European Commission (ERC-2024-POC, grant agreement no. 101213598, acronym SPINETRACER) and the Caixa Impulse program from La Caixa Foundation (CI23-10375). NA was supported by the Spanish National Plan for Scientific and Technical Research and Innovation— Ramon y Cajal (grant no. RYC2023-043851-I), the Consolidación Program of the Spanish State Research Agency (grant no. CNS2024-154900), and IKERBASQUE (grant no. RF/2023/006). MP, recipient of the AXA Chair, acknowledges the AXA Research Fund.

