## Supplementary Information for "Effect of carbon nanotubes in electroactive neuron and cardiomyocyte differentiation on conductive 3D printed scaffolds"

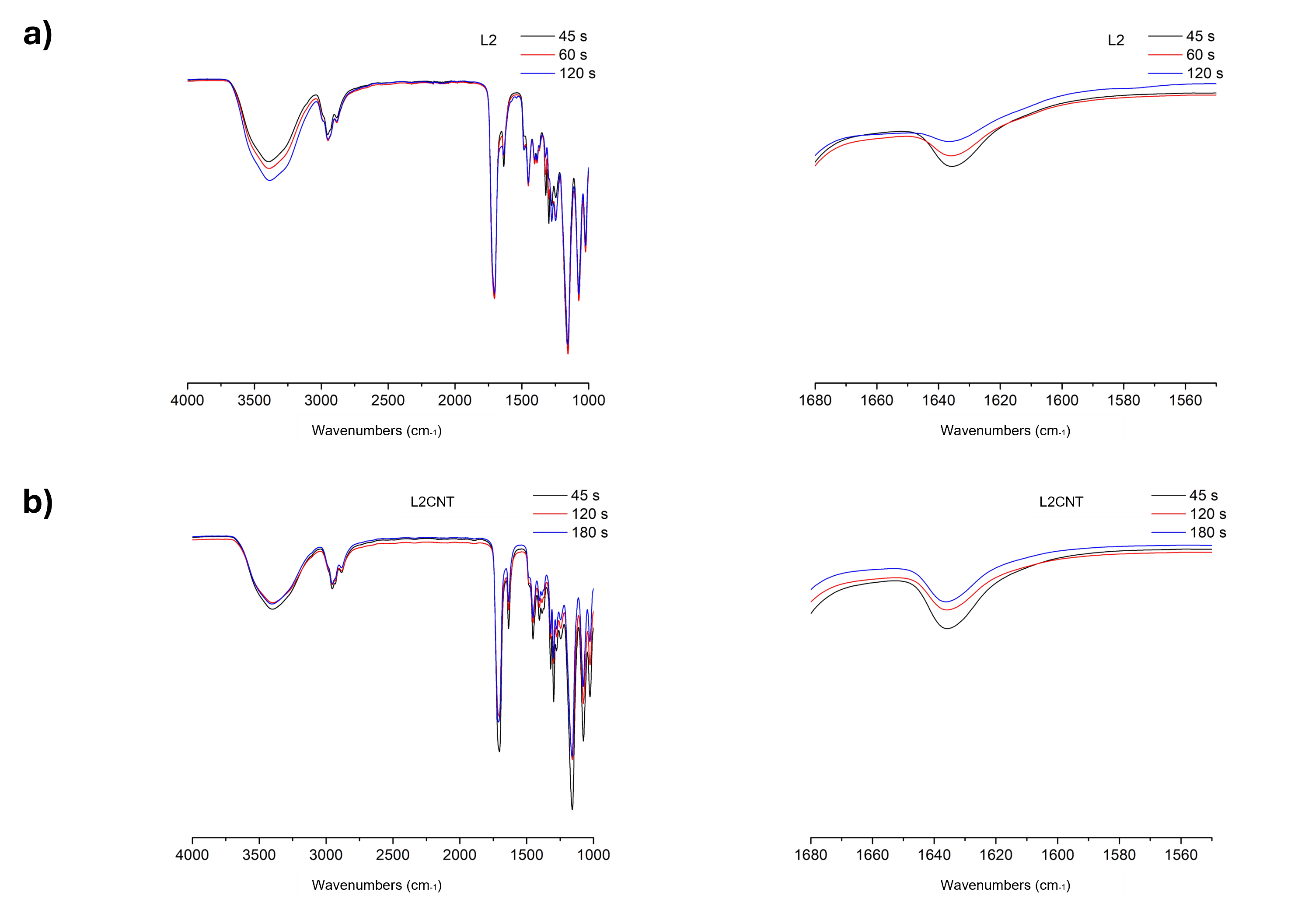
 **Figure S1.** FTIR and the corresponding zoom of a) L2 and b) L2CNT.

 
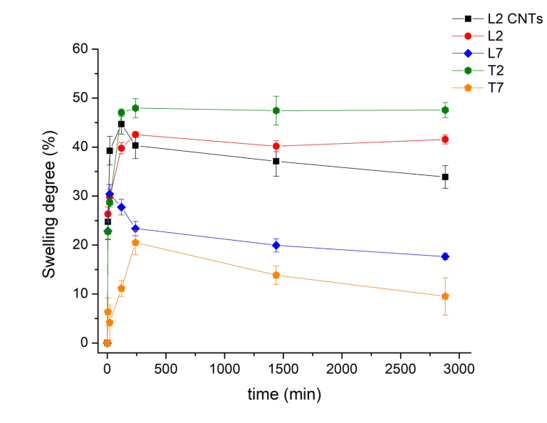
**Figure S2.** Swelling behavior of the scaffolds in PBS at 37°C.

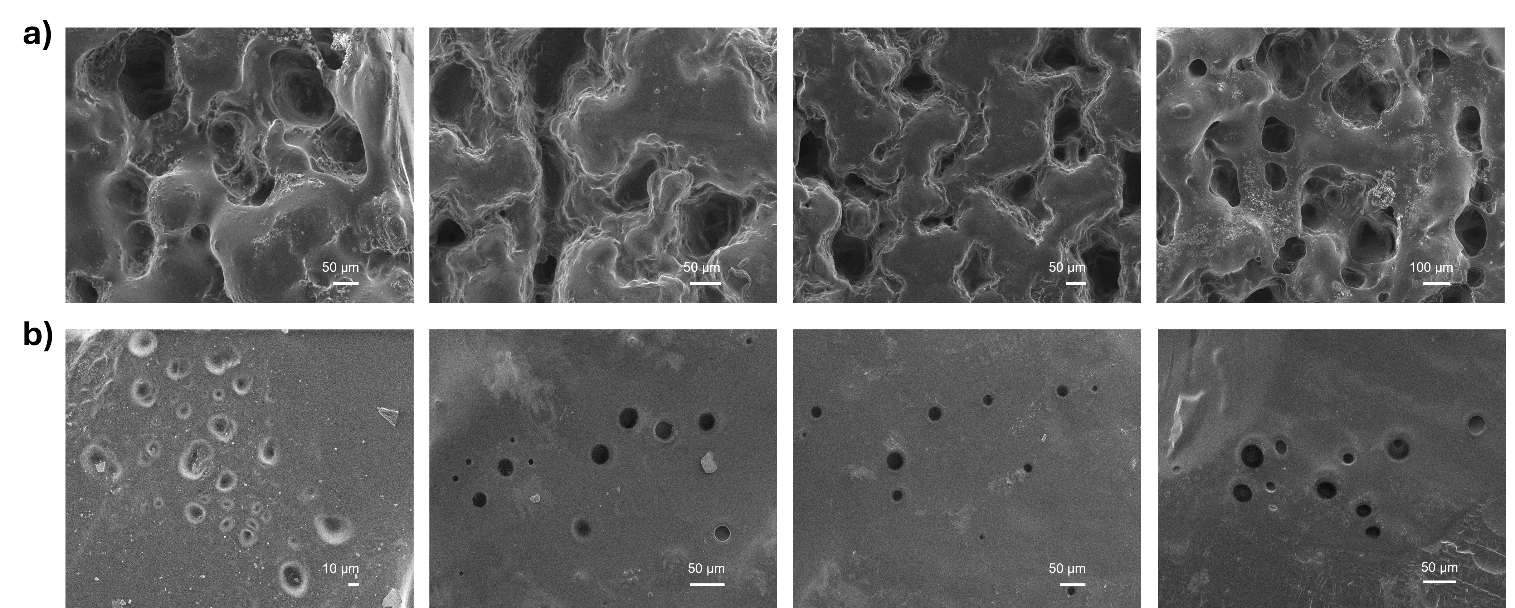

**Figure S3.** SEM images of the cross-sections acquired at an accelerating voltage of 2.50 kV for a) L2CNT and b) L2 scaffolds.

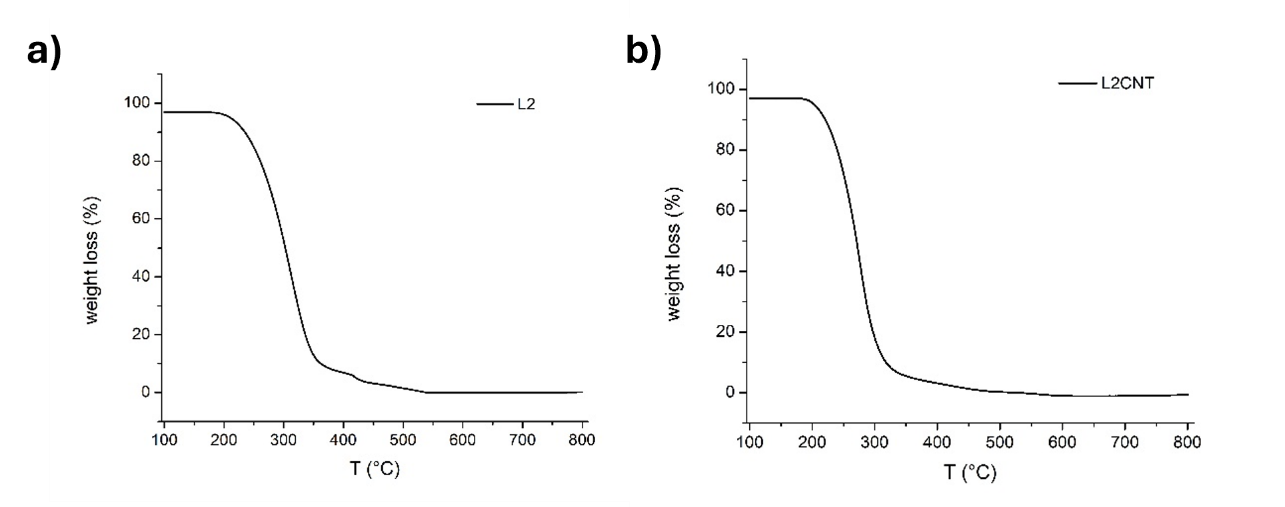
**Figure S4.** Thermogravimetric analysis results for a) L2 and b) L2CNT.

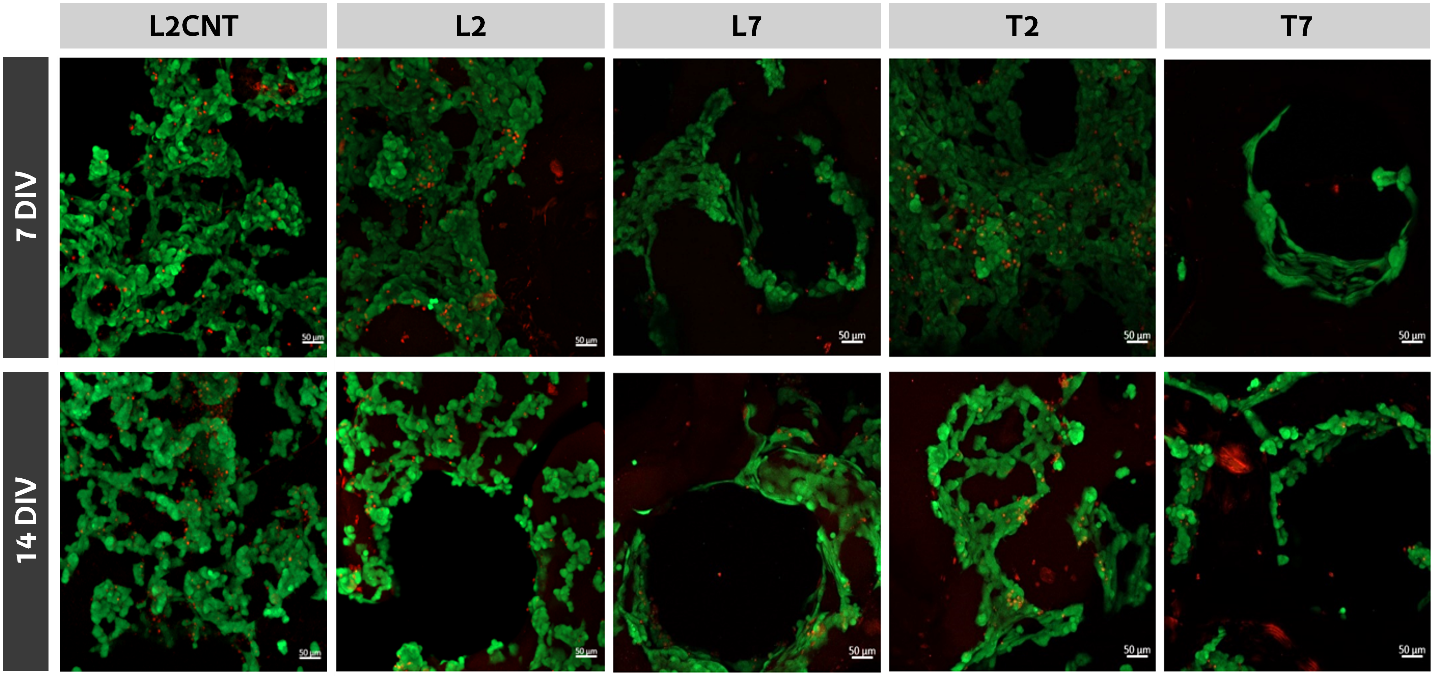

**Figure S5.** Live/Dead confocal images of the cells seeded on the scaffolds at 7 and 14 days in vitro. Live cells in green (CalceinAM, 488nm) and dead cells in red (BOBO-3, 570nm). Scale bars= 50µm.

**Table S1.** List of forward and reverse primer sequences used in RT-qPCR.

| **Target gene** | **Forward sequence** | **Reverse sequence** |
| --- | --- | --- |
| **GAPDH** | 5 ́- ACAGTTGCCATGTAGACC | 5 ́ - TTGAGCACGGGTACTTTA |
| **TUBB3** | 5’- TCATCTTTGCTCAGACTGG | 5’ - GTTTTCACACTCCTTCC |
| **NGN2** | 5’ - AAGATGTTCGTCAAATCCG | 5’ - CTCCTCCTCCTCTTCTTC |
| **ASCL1** | 5’- GAATGGACTTTGGAAGCAG | 5’- TTTTCTTTTCCTTTTCTCCCC |
| **SLC1A1** | 5’- CGAAAGAACCCTTTCCGATTTGC | 5’- GAAGGTGACAGGCAGTGTTGCT |
| **SOX2** | 5’- GCTACAGCATGATGCAGGACCA | 5’- TCTGCGAGCTGGTCATGGAGTT |
